# Thiazolidinediones are partially effective bitter blockers

**DOI:** 10.1101/2023.08.08.552460

**Authors:** Ha Nguyen, Cailu Lin, Ivona Sasimovich, Katherine Bell, Amy Huang, Emilia Leszkowicz, Nancy E. Rawson, Danielle R. Reed

## Abstract

**Purpose:** The bad bitter taste of some medicines is a barrier to overcoming non-compliance with medication use, especially life-saving drugs given to children and the elderly. Here we evaluated a new class of bitter blockers (thiazolidinediones; TZDs).

**Methods:** In this study, two TZDs were tested, rosiglitazone (ROSI) and a simpler form of TZD, using a high-potency sweetener as a positive control (neohesperidin dihydrochalcone, NHDC). We tested bitter-blocking effects using the bitter drugs tenofovir alafenamide fumarate (TAF), a treatment for HIV and hepatitis B infection, and praziquantel (PRAZ), a treatment for schistosomiasis, by conducting taste testing with two separate taste panels: a general panel (N=97, 20-23 yrs, 82.5% female, all Eastern European) and a genetically informative panel (N=158, including 68 twin pairs, 18-82 yrs, 76% female, 87% European ancestry). Participants rated the bitterness intensity of the solutions on a 100-point generalized visual analog scale.

**Findings:** Participants in both taste panels rated the bitter drugs TAF and PRAZ as less bitter on average when mixed with NHDC than when sampled alone. ROSI partially suppressed the bitterness of TAF and PRAZ, but effectiveness differed between the two panels: bitterness was significantly reduced for PRAZ but not TAF in the general panel and for TAF but not PRAZ in the genetically informative panel. ROSI was a more effective blocker than the other TZD.

**Implications:** These results suggest that TZDs are partially effective bitter blockers, suggesting other TZDs should be designed and tested with more drugs and on diverse populations to define which ones work best with which drugs and for whom. The discovery of bitter receptor blockers can improve compliance with medication use.

## Introduction

Non-compliance with medication use is a major source of outpatient treatment failure, and the bitter taste of medicines, especially crushed pills and liquid formulations given to children, is an important contributor^1,2^. Many unacceptably bitter medicines are life-saving drugs that must be administered to children too young and are used to treat common infectious diseases in resource-limited countries, such as tenofovir alafenamide fumarate (TAF) to treat HIV and hepatitis B infection and praziquantel (PRAZ) to treat schistosomiasis. Caregivers and health care providers often struggle with children when administering bitter medicines, and almost one-third of children with chronic conditions reported medicine refusal^3^. Techniques are available to mask the bitter taste of drugs by encapsulating the bitter active pharmaceutical ingredient into a solid form such as pills or tablets or to improve the poor palatability of liquid formulations by adding sweeteners and flavoring agents^4^. However, these approaches are limited to certain formulation options and dose requirements. Solid formulations can be problematic for children and patients who experience swallowing problems, and flavoring agents do not always mask the bitter taste adequately in liquid formulations^5^. Therefore, developing more effective approaches to block bitter taste is an increasing interest in both industry and academia.

An approach of interest is to block the bitter taste at the taste cell and receptor levels. Among at least 25 different type 2 taste receptors (TAS2Rs) involved in human bitter taste perception, most of them respond to multiple bitter-tasting ligands; likewise, bitter-tasting ligands can stimulate multiple TAS2Rs, resulting in a range of compounds that are perceived as bitter^6–8^. Blocking of bitter taste perception can be achieved by compounds that interact with taste receptors or the taste transduction pathway. Several compounds, including biomolecules^9,10^, flavanones^11,12^, and food-derived peptides and amino acid derivatives^13–15^, have been reported as bitter blockers for specific bitter receptors. However, their bitter blocking efficacy is limited, and a limited number of them have been tested in human participants for their efficacy and/or safety^5,14,16^.

One strategy to identify potential bitter blockers is to use cell lines expressing the bitter taste receptors for high-throughput biomolecular screening via calcium imaging^9^. However, any discovered or synthesized candidate antagonists need a risk assessment for their use in human participants. Therefore, as a first step toward achieving our goal to increase compliance with the treatment of common infectious and life-threatening diseases in resource-limited countries, in this study we screened drugs approved by the FDA, using bitter-response human taste-bud-derived epithelial cell culture platforms^17^ to identify bitter blockers for TAF. The thiazolidinedione (TZD) class of drugs was identified as potential bitter blockers. TZDs are commonly used in the treatment of type 2 diabetes, such as the FDA-approved drug rosiglitazone (ROSI; 5-(4-(2-(methyl(pyridin-2-yl)amino)ethoxy)benzyl)thiazolidine-2,4-dione)). We additionally tested TZDs for their blocking effects on PRAZ, another bitter drug commonly used for parasitic infections in resource-limited countries, to compare with effects on TAF.

While many medicines are bitter, they may not be bitter for everyone; similarly, the blocking efficacy may vary widely from person to person as well. Due to the heterologous expression of TAS2Rs, bitter perception and blocking effectiveness are specific for certain ligands that engage specific bitter receptors. The individual differences can partly be explained by genetic variation in bitter receptors, as well as other mediators such as saliva composition, age, psychiatric conditions, and previous experience (i.e., an adaptation to repeated exposure)^18–21^. The thyroid medication propylthiouracil (PROP) is well known for its difference in bitterness intensity due to genetic variation in the *TAS2R38* bitter receptor^19,22^. Quinine (malaria medication) and methimazole (thyroid medication) are other examples of drugs with individual differences in bitterness as a result of bitter receptor genotypes^22,23^. Thus, although the use of human taste-cell-based assays is a critical and relatively inexpensive method to understand and profile the bitter compounds and bitter blockers, the potential bitter blockers screened through cell lines may not block bitterness effectively in all human participants. For example, in recent work the terpenoid 6-methylflavone blocked TAS2R39 responses to TAF at both the cellular and the TAS2R receptor level; however, it did not work for 50% of the 16 human participants it was tested on, and it worked better for some people than for others^12^. Therefore, taste testing with human participants is necessary to identify new bitter blockers, and a genetically informative taste panel would be useful to examine origins of individual differences.

In this study, two TZDs, ROSI and a simpler TZD (see below), were tested with human participants and compared with the sweetener neohesperidin dihydrochalcone (NHDC) for their bitter suppression efficacy for TAF and PRAZ. NHDC, a semi-natural sweetener manufactured from neohesperidin, the parent flavanone obtained from bitter orange, has been reported to suppress bitterness^16,24,25^. We used both a general taste panel and a panel comprising mostly twin pairs, with greater diversity in age and race, to understand the effects of inherited variation on person-to-person differences in bitter blocker efficacy.

## Methods

### Drug and bitter blocker selection and sample preparation

The characteristics and concentrations of drugs, bitter blockers, and diluents used in this study are presented in **Table 1**. We used PROP as a control drug because of its well-known bitterness intensity and genetic effects^19,22^. Blocking efficacy of a simpler thiazolidine-like analog compound, 3-(hydroxymethyl)-5-(4-methylbenzylidene)-2-thioxo-1,3-thiazolidin-4-one (referred to hereafter as simpler TZD), was used to compare with ROSI.

**Table 1.** Characteristics of drugs, bitter blockers, and diluents used in this study.

| Compound (Abbreviation) | Usage | PubChem CID | Taste quality | Diluent | Sample concentration (μM) | Supplier (Catalog no.) |
| --- | --- | --- | --- | --- | --- | --- |
| **Bitter drugs** |  |  |  |  |  |  |
| Tenofovir alafenamide fumarate (TAF) | HIV medication | 9574768 | Bitter | Deionized water | 200 | Nanjing Bilat Chemical Industrial Co (202138-50-9) |
| Praziquantel (PRAZ) | Schistosomiasis medication | 4891 | Bitter | 3% ethanol | 300 | Sigma Aldrich (P4668-5G) |
| Propylthiouracil (PROP) | Thyroid medication | 657298 | Bitter | Deionized water | 560 | Pfaltz & Bauer, Inc. (P28100) |
| **Bitter blockers** |  |  |  |  |  |  |
| Rosiglitazone (ROSI) | Type 2 diabetes medication | 77999 | — | 2% Tween 80 | 140 | TCI Chemicals (R0106-1G) |
| 3-(Hydroxymethyl)-5-(4-methylbenzylidene)-2-thioxo-1,3-thiazolidin-4-one (“simpler TZD”) | — | — | — | 2% Tween 80 | 140 | Chembridge Corporation (5404644) |
| Neohesperidin dihydrochalcone (NHDC) | Food additive (sweetener) | 30231 | Sweet | Deionized water | 130 | TCI Chemicals (N0675-5G) |

We conducted tests with a pilot taste panel (N=13, 20-49 yrs, 46% female) to identify the concentration of each drug that gave mid-range intensity ratings of bitterness and a concentration of each bitter blocker that showed a blocking effect on the bitter drug. The amount of ROSI was 0.5 mg when applying 10 ml of 140 µM ROSI solution as a pre-rinse (sip and spit), an amount much lower than the FDA-approved dosage for medication (commonly starting at 4 mg/day), which ensures minimal risks of taking this drug as a blocker. The drugs and bitter blockers were purchased from the vendors listed in **Table 1**. Participants received 10 mL solution samples at ambient temperature in 30 mL white plastic bottles with screw caps.

### Human sensory testing

We conducted tests with two separate taste panels. First, with the general panel we tested the suppression efficacy of ROSI and NHDC when applying them either mixed with the drug or as a pre-rinse agent. We retested the bitter blockers/maskers with the genetically informative panel to understand effects of inherited variations on person-to-person differences in bitter blocker efficacy. The simpler TZD became available after sensory tests with the general panel were complete, so it was tested only with the genetically informative panel. The potency of the blocker/masker was tested on the two bitter drugs TAF and PRAZ. The general panel data showed that mean bitterness intensity ratings were significantly lower with the mixtures than with the pre-rinses with NHDC with TAF and ROSI with PRAZ, but not in other samples (**Table S1**). Thus, we chose to use the easier method in the genetically informative panel, applying the bitter blocker/masker only as a mixture with the bitter medicine.

For each panel, participants completed a 30-minute tasting session to evaluate the overall bitterness intensity of three warm-up solutions (distilled water, ROSI, and TAF), three drug solutions (TAF and PRAZ as bitter test drugs, and PROP as a control drug to check data reliability and validity according to *TAS2R38* receptor genotype), and mixtures/pairs of the blocker/masker with the drug (four mixtures and four pairs in the general panel, and six mixtures in the genetically informative panel).

The concentration of each drug or bitter blocker/masker was the same for both taste panels and across warm-up, single, and paired solutions within each session (**Table 1**). To assess perceived bitterness intensity, we used a generalized visual analog scale (gVAS) to allow valid comparisons across different participant groups with different levels of sensory experience^26,27^. Participants were asked to rate the bitterness intensity of the sample on a horizontal line scale ranging from 0 = “no bitterness” to 100 = “strongest imaginable bitterness”. Participants were first provided with verbal and written instructions on how to use the scale and then started with three warm-up samples to familiarize themselves with bitter perception and scale usage. The data for warm-up samples were used for data quality checks. Warm-up samples were presented in a fixed order across participants: distilled water (control sample), ROSI, and TAF, respectively. ROSI was possibly a low-bitter compound and was included to test its bitterness. TAF was used as a highly bitter reference and a replicate to check test-retest reliability within each tasting session. The sample orders of single or paired solutions were balanced among participants using Williams’ Latin square designs. All samples were blinded with random 3-digit codes.

For each evaluation, participants were instructed to place a nose clip on, put the whole sample in their mouth, swish it for five seconds, and spit it out. They rated the bitterness intensity while the sample was in their mouth. They were asked to cleanse their palates with drinking water and unsalted crackers (if needed) and wait until perceiving no sensation in their mouths (at least 30 sec) before evaluating the next sample. For paired solutions, they rated the bitterness intensity of the second sample without rinsing the palate between the two samples. Before participants started the taste test, they were asked to provide a saliva sample and complete a survey regarding their demographics (age, gender) and health. DNA purification and genotyping were performed on the saliva sample to identify their *TAS2R38* genotypes^28^. For the general panel the taste test and survey were conducted in person using paper ballots, and the genetically informative panel used RedJade software (RedJade Sensory Solutions, Martinez, CA).

### Participants

The general panel included 97 volunteer students (20–23 yrs, 82.5% female, all of Eastern European ancestry) at the University of Gdańsk (Gdańsk, Poland) (**Table 2**). The genetically informative panel comprised 158 participants, including 68 twin pairs and 22 singletons, recruited at the 2022 Twins Days Festival (Twinsburg, Ohio, USA). The genetically informative panel participants were more diverse in age and race (18–82 yrs, 76% female, 87% White or Caucasian). The exclusion criteria included pregnancy and age under 18 yrs. A subset of the genetically informative panel (n=20, 20–57 yrs, 75% female) was tested twice, on different days, to assess the test-retest reliability of their responses.

**Table 2.** Participant characteristics.

|  | Test panel | Retest panel |
| --- | --- | --- |
| **General panel** |  |  |
| Total participants, N (%) | 97 (100.0) | — |
| Age, median [interquartile range] | 22 [21, 23] | — |
| Gender, N (%) |  |  |
| Male | 17 (17.5) | — |
| Female | 80 (82.5) | — |
| Ethnicity, N (%) |  |  |
| White or Caucasian | 97 (100.0) | — |
| **Genetically informative panel** |  |  |
| Total participants, N (%) | 158 (100.0) | 20 (100.0) |
| Age, median [interquartile range] | 34 [24, 46] | 36 [34, 45] |
| Gender, N (%) |  |  |
| Male | 38 (24.0) | 5 (25.0) |
| Female | 120 (76.0) | 15 (75.0) |
| Ethnicity, N (%) |  |  |
| White or Caucasian | 137 (87.0) | 18 (90.0) |
| Hispanic or Latino | 2 (1.3) | 2 (10.0) |
| Black or African American | 15 (9.5) | — |
| Other or mixed ancestry | 4 (2.5) | — |
| Twin status |  |  |
| Monozygotic | 114 (72.2) | 16 (80.0) |
| Dizygotic | 18 (11.4) | 2 (10.0) |
| Singleton | 22 (13.9) | 2 (10.0) |
| Unknown | 4 (2.5) | — |

All study procedures were approved by the Institutional Review Board at the University of Pennsylvania (protocol #701426, Genetic Studies of Taste and Oral Perception) and the Bioethical Committee by the Regional Medical Chamber in Gdansk, Poland. The sip-and-spit procedures for the active pharmaceutical ingredients of FDA- and WHO-approved drugs and common sensory compounds confer no more risk than from a routine medical visit. All participants completed informed consent before participating in the sensory test. Participants in the genetically informative panel received a gift card in acknowledgment of their time and contribution to the study; the general panel received university course credit.

### Data analysis

Raw ratings (from 0 to 100) for all participants in each taste panel were used to assess the data reliability and individual differences in the perceived bitterness intensity of the drugs with or without the bitter blocker. The results are presented in pirate plots with medians, minimum and maximum values, and interquartile ranges (IQR: 25th to 75th percentile) to compare rating distributions (**Figure 1**). It is not possible to evaluate the potency of a bitter blocker for participants who are insensitive to the bitter taste of a particular stimulus. Therefore, we removed the data for participants who rated both TAF and PRAZ as not or slightly bitter (with ratings ≤ 25 on the 100-point gVAS), which is 55 participants in the general panel and 51 participants in the genetically informative panel. We also removed all data for eight participants in the genetically informative panel who rated the water warm-up sample as moderate to extreme bitterness intensity (>25 on the gVAS). For these two subgroups after data removing (N=42 in the general panel, and N=99 in the genetically informative panel), a bitter suppression score was calculated by subtracting the bitterness intensity rating of the mixture from that of the drug, presented in percentage bitterness compared to the drug alone (**Table 3**); thus, larger suppression scores mean more bitter suppression. For comparing raw ratings (**Table S1**) or bitter suppression scores (**Table 3**) between the drugs, the drugs mixed with the blockers, and/or the drugs with the blockers applied as a pre-rinse, linear mixed-effects models were performed, with samples as a fixed effect and participants as a random effect.

**Figure 1.**
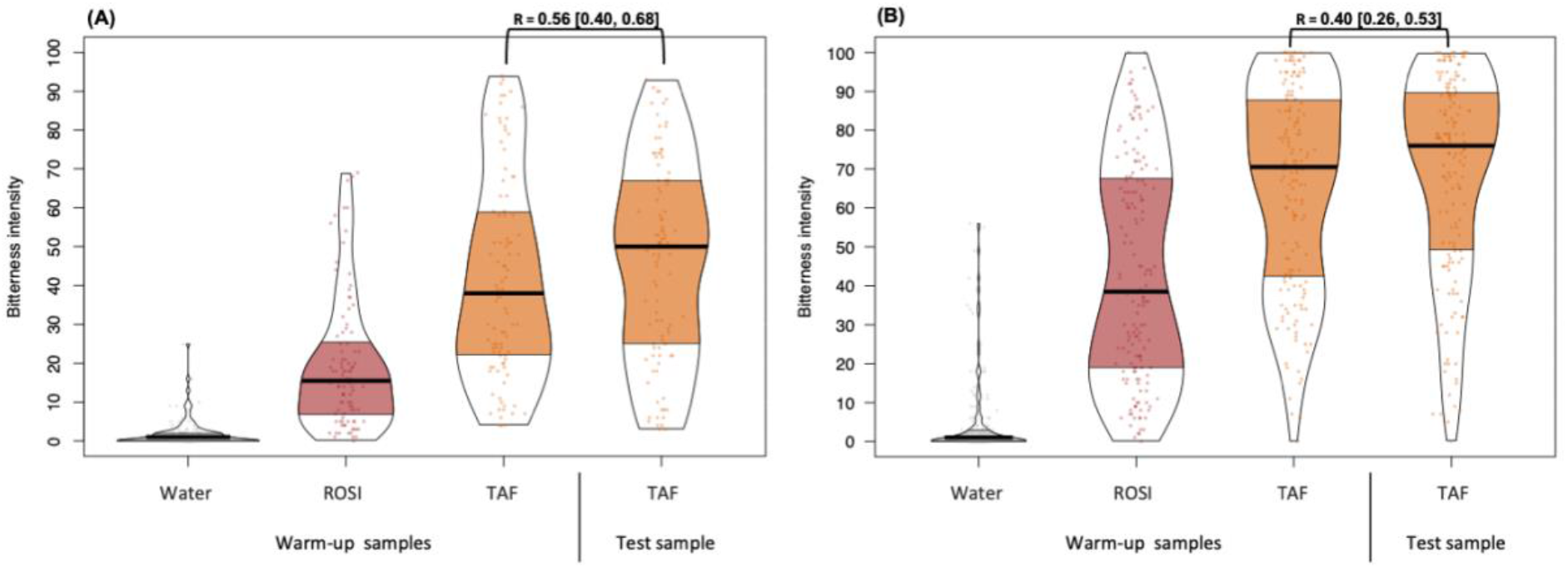
Reliability of bitterness ratings in the general panel (A) and the genetically informative panel (B). Participants rated the bitterness intensity of solutions using a 100-point generalized visual analog scale. Points represent individual data (jittered horizontally); the central lines depict the median, the shapes reflect the density of the distribution, and the shaded areas show interquartile ranges. R [CI 95%] is Spearman’s correlation between the warm-up and test TAF ratings. For abbreviations, see Table 1.

**Table 3.** Bitter suppression of TAF and PRAZ with blockers for participants who perceived drugs as bittera.

| Blocker | Ligand | Suppression (%) |  | Statistical grouping<sup>b</sup> |
| --- | --- | --- | --- | --- |
|  |  | Mean | [CI 95%] |  |
| General panel (N=42) |  |  |  |  |
| NHDC | TAF | 42.9 | [30.3, 55.6] | AB |
|  | PRAZ | 34.3 | [21.6, 46.9] | B |
| ROSI | TAF | 7.2 | [-5.4, 19.8] | C |
|  | PRAZ | 58.8 | [46.1, 71.4] | A |
| Genetically informative panel (N=99) |  |  |  |  |
| NHDC | TAF | 30.5 | [20.3, 40.6] | AB |
|  | PRAZ | 45.0 | [34.9, 55.2] | A |
| ROSI | TAF | 23.4 | [13.3, 33.6] | BC |
|  | PRAZ | 8.9 | [-1.3, 19.0] | CD |
| Simpler TZD | TAF | -0.5 | [-10.6, 9.7] | D |
|  | PRAZ | 5.7 | [-4.5, 15.8] | D |

For abbreviations, see Table 1.
<sup>a</sup>: Participants who rated TAF and PRAZ > 25 on the 100-point generalized visual analog scale.
<sup>b</sup>: Different letters show significant differences in mean suppression (%) between samples within one taste panel. All results reflect the bitter blocker given as a mixture with the bitter drug.

The genetically informative panel comprised 68 twin pairs, including 57 monozygotic (MZ) and 9 dizygotic (DZ) twin pairs, 2 unknown twin pairs, and 22 other individuals (**Table 2**). There were significant correlations in ratings of six of nine test solutions between individuals within MZ twin pairs (**Figure 2**), and the number of DZ twin pairs was too small to detect any correlations (**Figure S1**). Thus, to account for the twin correlation, the identification of twin status (pedigree #) was included as a random effect in the ANOVA models. Tukey’s HSD (honestly significant difference) test was used for post-hoc pairwise comparisons of sample means^29^. The results are presented in means and 95% confidence intervals (CI 95%) in **Table 4**, **Table S1**, and **Table S2**.

**Figure 2.**
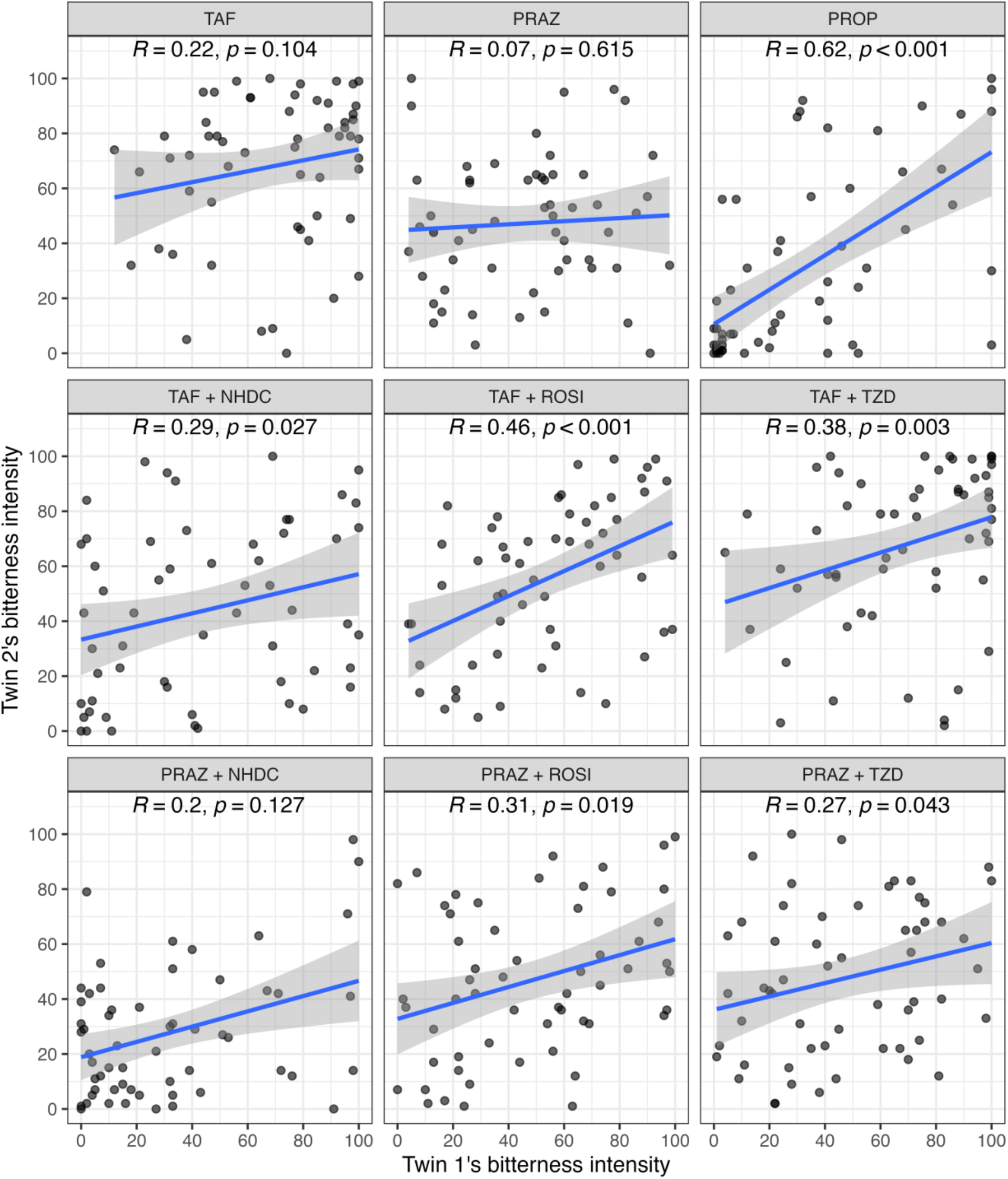
Spearman’s correlation in bitterness intensity ratings between twin 1 and twin 2 for MZ twin pairs in the genetically informative panel (N=57). For abbreviations, see Table 1.

**Table 4.** Bitter suppression for three clusters with distinct suppression score profiles identified in the genetically informative panel by hierarchical clustering on principal components.

| Blocker | Ligand | TZD Effect Cluster<sup>a</sup> |  |  |  |  |  |
| --- | --- | --- | --- | --- | --- | --- | --- |
|  |  | Ineffective |  | Selectively effective |  | Effective |  |
|  |  | (N=21) |  | (N=37) |  | (N=41) |  |
| NHDC | TAF | 35.4 | [18.0, 52.0] B | -5.4 | [-16.0, 5.0] A | 59.6 | [50.0, 69.0] C |
|  | PRAZ | 41.1 | [17.0, 65.0] B | 11.8 | [-6.6, 30.0] A | 76.4 | [70.0, 83.0] C |
| ROSI | TAF | -3.2 | [-20.0, 14.0] A | 14.7 | [1.9, 28.0] A | 44.3 | [35.0, 53.0] B |
|  | PRAZ | -64.3 | [-94.0, -34.0] A | 18.7 | [6.6, 31.0] B | 36.8 | [25.0, 48.0] B |
| Simpler TZD | TAF | -18.3 | [-46.0, 9.1] A | -4.2 | [-19.0, 10.0] AB | 11.3 | [0.4, 22.0] B |
|  | PRAZ | -72.6 | [-92.0, -53.0] A | 16.8 | [3.6, 30.0] B | 35.0 | [23.0, 47.0] B |

<sup>a</sup>: Different letters show significant differences in mean suppression (%) between clusters (column) within each sample (row). All results reflect the bitter blocker given as a mixture with the bitter drug.

For the genetically informative panel we performed principal component analysis and hierarchical clustering on principal components analysis on suppression scores, with Euclidean distances and Ward’s clustering method^30^, to identify possible participant clusters and understand individual differences in the bitter blocker efficacy. All statistical analyses were performed using R 4.2.1^31^ and RStudio 2022.07.2.576^32^, with significance set at p-value ≤ 0.05.

The R scripts and data are available on GitHub (*the link will be available when this paper is published*).

## Results

### Participant characteristics, overall perceived bitterness, and data reliability

Most of the genetically informative panel participants were White or Caucasian (87%), although they were more diverse in age and ethnicity than the general panel participants (**Table 2**). In the genetically informative panel, the demographic characteristics were similar for the first session (the test panel, N=158) and the second session (the retest panel, N=20). Among 99 participants in the genetically informative panel who perceived the drugs as bitter (>25 on the 100-point gVAS), the association between age and the suppression score was statistically significant only for TAF + TZD (R-Spearman = 0.32, p-value = 0.001); however, no significant correlations between age, race, or sex and bitterness ratings or other blocker efficacy were identified.

There was a large variation in the perceived bitterness intensity of the drugs among participants, with greater variation in the genetically informative panel than in the general panel. Drugs and bitter blockers were not bitter for some participants but moderately to highly bitter to others (see, e.g., bitter rating distributions for TAF and ROSI in **Figure 2**). Among 97 participants in the general panel, about 43% (N=42) rated TAF and PRAZ alone as bitter (>25 on the 100-point gVAS), as did about 63% (N=99 of 158) in the genetically informative panel. The two participant groups provided different bitterness ratings, with greater means in the genetically informative panel (**Table S2**) than in the general panel (**Table S1**); for example, the mean TAF bitterness ratings were 68.1 and 46.5, respectively.

The data reliability of the two panels was confirmed by the ability to distinguish the bitterness of water, slightly bitter, and highly bitter solutions, by significant correlations between the warm-up and test samples within one tasting session (**Figure 2**), and by high test-retest reliability across two tasting sessions in the genetically informative panel (**Figure S2**). In the general panel, all participants rated the water sample ≤25 on the bitterness intensity scale and provided a significant correlation between warm-up and test TAF ratings (R = 0.56, p-value ≤ 0.05). In the genetically informative panel, Spearman’s correlation between the warm-up and test TAF samples was 0.40 (p-value ≤ 0.05), which increased to 0.55 after removing the data of the eight participants who rated the bitterness of water as >25 on the gVAS. This confirms the need to remove those unreliable data points for further statistical analyses to compare the efficacy of different bitter blockers. Although the sample size of the retest panel was small (N=20), the genetically informative panel showed test-retest reliability (R = 0.42–0.60, p-value ≤ 0.05 for all test samples; **Figure S2**).

### Genetic effects on the bitterness intensity of medicines and bitter blocker efficacy

Genetic differences in bitter receptors can explain individual differences in bitter perception^19^. In the case of PROP bitterness, variants in the *TAS2R38* bitter receptor gene result in differences in bitter perception: AVI is the bitter insensitive haplotype, and PAV is the bitter sensitive haplotype. For both panels, PROP bitterness intensity was significantly lower in participants with AVI/AVI diplotype (non-tasters) than in those with AVI/PAV diplotype (medium tasters) or PAV/PAV diplotype (super-tasters) (**Figure 3**). This expected relationship confirmed the reliability and validity of the data. There were no significant differences in TAF or PRAZ bitterness intensity across the PROP sensitivity groups. Moreover, individual ratings of the drugs (TAF, PRAZ, and PROP) were moderately correlated (R=0.17–0.31, p-value ≤ 0.05), indicating a shared determinant of the person-to-person differences in perceived bitterness of these drugs.

**Figure 3.**
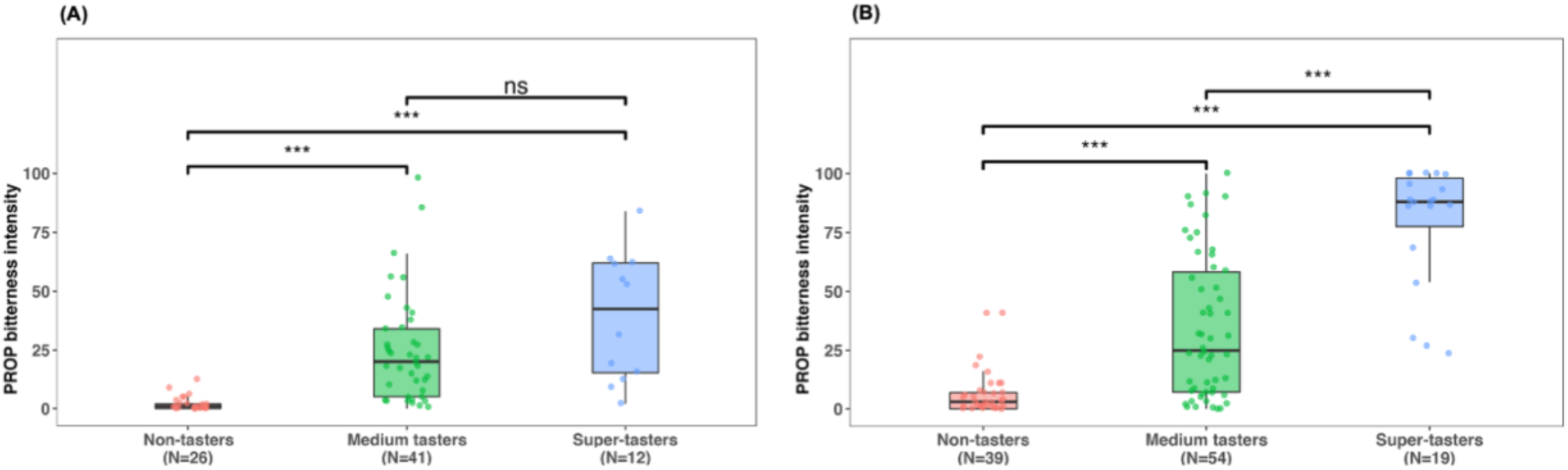
Propylthiouracil (PROP) bitterness intensity by PROP sensitivity group in the general panel (A) and the genetically informative panel (B): non-tasters (insensitive to PROP), medium tasters (intermediate sensitivity to PROP), and super-tasters (highly sensitive to PROP). Points represent individual data (jittered horizontally); the central lines depict the median, and the shaded areas show interquartile ranges. *** *p* ≤ 0.001; ns, not significant.

In the genetically informative panel, the correlation in bitter ratings between individuals within MZ twin pairs (N=57) was significant for PROP (R=0.62); for TAF mixed with NHDC (R=0.29), ROSI (R=0.46), or the simpler TZD (R=0.38); and for PRAZ mixed with ROSI (R=0.31) or the simpler TZD (R=0.27) (**Figure 1**); among the small number of DZ twin pairs (N=9) no significant correlations were identified. No significant correlations in suppression scores were identified among a small number of MZ twin pairs (N=23) or DZ twin pairs (n=3) who rated TAF and PRAZ alone as bitter (data not reported). The high MZ correlation for PROP again confirmed the genetic effect on PROP bitterness perception. The moderate MZ correlations for most of the drug and blocker mixtures suggest that there is inherited variation in the bitter perception of these drugs and potentially in bitter blocker efficacy.

### NHDC partially suppressed the bitterness of TAF and PRAZ

As expected, the addition of NHDC reduced the bitterness of both TAF and PRAZ for both panels. For participants who rated TAF and PRAZ as bitter (>25 on the 100-point gVAS), moderate suppression (averaging 30.5–45%) was observed for both TAF and PRAZ plus NHDC in both panels. The average suppression scores were not significantly different between drugs tested within each panel: 42.9% and 34.3% for TAF and PRAZ, respectively, in the general panel, and 30.5% and 45% in the genetically informative panel (**Table 3**). The raw ratings for all participants showed similar suppression for NHDC: participants in both panels rated the bitter drugs TAF and PRAZ as less bitter on average when mixed with NHDC than when sampled alone (**Table S1**, **Table S2**).

The hierarchical clustering on principal components analysis for the genetically informative panel revealed three clusters of participants with distinct suppression score profiles representative of the efficacy of the different TZDs: ineffective (TZDs did not work for both TAF and PRAZ), selectively effective (certain TZDs worked for certain drugs), and effective (TZDs worked for both TAF and PRAZ) (**Table 4**). The suppression with NHDC was high for the effective TZD cluster (59.6% for TAF, 76.4% for PRAZ, N=41), moderate for the ineffective TZD cluster (35.4% for TAF, 41.1% for PRAZ, N=21), but not significant for the selectively effective TZD cluster (N=37) (**Table 4**).

### The blocking efficacy of thiazolidinediones differed between participant groups

#### ROSI and the simpler TZD had low to moderate bitter suppression for certain drugs for certain participants

ROSI partially suppressed the bitterness of TAF and PRAZ, but with different results for the two panels. In the general panel ROSI significantly reduced the bitterness of PRAZ but not TAF, whereas in the genetically informative panel it reduced the bitterness of TAF but not PRAZ (**Table 3**, **Table S1**, **Table S2**). Among the effective suppressions (statistically significant bitterness reductions), the potency of ROSI and NHDC were not significantly different in the general panel (58.8% for PRAZ + ROSI vs. 34.3% for PRAZ + NHDC) or in the genetically informative panel (23.4% for TAF + ROSI vs. 30.5% for TAF + NHDC).

The simpler TZD worked on a subgroup of participants in the genetically informative panel. The mean suppression score for the simpler TZD mixed with TAF or PRAZ was not significantly different from zero for the whole panel (**Table 3**), but for the effective TZD cluster (N=41) it was an effective bitter blocker of TAF/PRAZ, although its efficacy was smaller than that of NHDC (**Table 4**). In the selectively effective TZD cluster (N=37) the simpler TZD was an effective bitter blocker for PRAZ but not for TAF. In the ineffective TZD cluster (N=21) neither ROSI nor the simpler TZD worked; in the other two clusters ROSI was a low to moderate effective blocker of TAF and PRAZ.

#### There were large individual differences in suppression efficacy

Among 99 participants in the genetically informative panel who perceived the drugs as bitter, about 16% (N=16) reported nearly full bitter suppression for TAF + NHDC (suppression scores > 75%), and 34% reported nearly full suppression for PRAZ + NHDC, compared to about 11% for TAF or PRAZ + ROSI, 8% for PRAZ + TZD, and 2% for TAF + TZD. Additionally, about 41% of participants (the effective TZD cluster, N=41; **Table 4**) reported greater bitter suppression of the drugs with the blockers than other participants in all cases, with high average suppression (>50%) for TAF or PRAZ + NHDC, moderate average suppressions (30–50%) for TAF or PRAZ + ROSI and PRAZ + TZD, and a low average suppression (<30%) for TAF + TZD. Noticeably, in the ineffective TZD cluster the addition of ROSI or the simpler TZD to the PRAZ solution increased the bitterness of the solution, causing significantly negative suppression in those participants. However, there were no significant associations between the three clusters and gender, age, or ethnicity.

## Discussion

TZDs are synthetic peroxisome proliferator-activated receptor gamma (PPARγ) ligands that enhance insulin secretory response and are used in the treatment of type 2 diabetes^33^. To our knowledge, there are no previous reports on the bitter-blocking effect of any TZDs. Using a screening platform, we identified TZDs that can block responses to the bitter drug TAF. However, blocking TAF-response receptors in cell culture may not reflect the suppression of bitter taste in taste testing^6^. In a recent study, a similar approach identified 6-methylflavone as a blocker of TAF bitterness. However, the bitter suppression was not statistically significant in a taste panel (N=16), although 50% of participants showed a reduction in perceived bitterness^12^. In this study, we conducted taste testing with the two TZDs, ROSI and the simpler TZD, plus NHDC as a positive control, for bitter suppression effect with the bitter drugs TAF and PRAZ. The TZDs were partially effective bitter blockers not only for TAF as identified by the platform but also for PRAZ included as a comparison, although there were person-to-person and panel-to-panel differences in suppression efficacy. This project builds on *in vitro* evidence to support the efficacy of both TZDs, but neither drug was completely effective at blocking bitterness in all participants, but medicinal chemistry modification to improve potency and efficacy is warranted; such effort may allow us to find a better analog(s) that antagonize multiple TAS2Rs.

The receptor-ligand relationship can be extremely specific; thus, the blocking application is specific for certain ligands that engage specific bitter receptors. A single universal blocker that acts by blocking each and every bitter receptor is unlikely. Additionally, small changes in receptor structure due to person-to-person differences can cause large variations among participants in the perception of bitter intensity of the same drug, and for bitter suppression with the same blocker^12,18–21^. Both ROSI and the simpler TZD worked as partially effective bitter blockers. However, blockers themselves can be bitter; e.g., the sweetener cyclamate suppresses the responses of the bitter receptors to sweetener saccharin, and vice versa^34^.

ROSI was perceived as slightly bitter by people in the general panel, who were of Eastern European ancestry, and moderately bitter on average by participants in the genetically informative panel, who were of more diverse ancestry. The simpler TZD was also perceived as slightly to moderately bitter by pilot testing. For participants on whom the TZDs were ineffective, the bitterness of ROSI and the simpler TZD might cause a significant increase in the bitterness intensity of PRAZ, which by itself was rated as moderately bitter. This result suggests that TZDs do not work for those participants. A mixture containing TZDs and antagonists of other bitter receptors would likely decrease the drug bitterness for a larger, heterogeneous population.

Inherited variation can partially explain the person-to-person differences in medicine bitterness and blocker efficacy. The results of *TAS2R38* genotypes and the twin correlations in the genetically informative panel helped to partially reveal the genetic origin of medicine bitterness and blocker efficacy. A future twin study with a larger sample size can quantify the heritability of blocker efficacy. The bitterness of TAF and PRAZ was not significantly correlated with *TAS2R38* receptor genes; thus, genotyping of bitter receptors may help us understand the genetic variation in the bitter perception of TAF and PRAZ and the blocking efficacy of TZDs. It is reported that TAF can activate TAS2R39, TAS2R1, TAS2R8, and TAS2R14^12^, but there is no available data on the associated receptors for PRAZ. Testing the blocking effect of TZDs on TAF and PRAZ with receptor-based heterologous expression assays^12^ will help us understand which receptors TZDs worked on as bitter blockers. Also, there may be other sources of genetic variation in bitter blocking beyond the bitter receptors themselves, such as the saliva components, which affect fungiform papillae density and maintenance^35^.

Although we compared the blocking effects using both pre-rinse and mixture procedures, their results were similar for half of drug-blocker pairs, so we chose the least complicated procedure. This observation suggests that the practical application of these bitter blockers can be either a solution/emulsion to be used with the bitter drug as a pre-rinse or mixed with the drug in liquid formulations. The blocker solution can be sold separately from the drug packages and customized for specific populations and drugs as needed. Bitter blockers can act independently of the co-administered compounds; thus, this approach can reduce product costs when the compound is delivered separately from the drug packages. The most likely end-user will be patients who are struggling with medicine administration due to the unacceptable bitter taste, for example, children and the elderly taking liquid formulations.

One limitation of this study is that taste tests were conducted in a class setting (the general panel) or at an outside event (the genetically informative panel), with less controlled environments than in a standard sensory laboratory. However, we included control samples and in-session test-retest samples in the experiment design to check data reliability and remove unreliable data for further analysis. We also checked the data reliability and validity through the expected relationship between the taste genotypes and phenotypes (i.e., PROP bitterness and *TAS2R38* diplotype). The results suggest that TZDs are partially effective bitter blockers.

However, perhaps due to the ancestry differences in the tested populations, the blocking effect of the two TZDs differed between the two panels. Bitter taste perception widely differs from person to person and across populations worldwide. The large variation in bitter suppression can be explained by the differences in participant genetics, demographics, and previous experience. Therefore, other TZDs should be tested with more drugs and on different participant groups to define which ones work best with which drugs.

## Conclusions

TZDs are potentially valuable as bitter blockers because they are (a) partially effective in most people and (b) part of a class of drugs approved worldwide for the treatment of diabetes (and therefore face a lower regulatory burden for use in medicines). Our findings of different responses from panelists from different parts of the world with different ancestries indicate person-to-person differences in bitter blocker efficacy. Thus, having more blockers to choose among will be necessary to fully suppress the bitterness of medicines for a broad range of populations and ancestries. Adding a mixture of bitter blockers to medicines, in conjunction with sweeteners and flavorings, may help attain a low- to zero-bitterness standard for the most noxious medicines.

## Acknowledgments

We acknowledge AeroNeph Therapeutics, Inc. (legacy DiscoveryBioMed, Inc.) and Drs. John Streiff and Erik Schwiebert from which and from their efforts, the simpler TZD-like drug fragment was discovered using bitter-responsive cell-based assays. We thank Yang Hu and Shanice James for their assistance in the sample preparation for sensory testing. We thank students, especially Joanna Pasztelan, from the Faculty of Biology at the University of Gdańsk, for their assistance in data collection in Gdańsk (Poland). We thank Charles Arayata, Ahmed Barakat, May Cheung, Lauren Colquitt, Daniel Doran, Justin Doran, Riley Herriman, Sarah Marks, Robert Pellegrino, and Aurora Toskala for their assistance in data collection at the 2022 Twins Days Festival (Twinsburg, Ohio, USA).

## Author declaration of individual contribution

Nguyen, Lin, Rawson, and Reed participated in research design. Nguyen and Sasimovich developed experimental protocols. Bell and Huang prepared testing samples. Nguyen, Sasimovich, Bell, Huang, Leszkowicz, and Reed collected data. Nguyen and Lin performed data analysis. Nguyen and Reed wrote the manuscript.

