## Supplemental tables and figures for "Thiazolidinediones are partially effective bitter blockers"

**Supplemental Materials**

**Table S1**. Bitterness intensity of drugs and the mixtures or pre-rinses with bitter blockers rated by the general panel (N=97) on a 100-point generalized visual analog scale.

| Blocker | Ligand | Sample | Bitterness intensity Mean [CI 95%] | Statistical grouping[a] |
|---|---|---|---|---|
| None | TAF | Single | 46.5 [41.5, 51.5] | A |
| | PRAZ | Single | 31.8 [26.7, 36.8] | BC |
| | PROP | Single | 19.5 [14.5, 24.5] | EF |
| NHDC | TAF | Mixture | 25.6 [20.6, 30.6] | CDE |
| | PRAZ | Mixture | 22.4 [17.4, 27.4] | DEF |
| ROSI | TAF | Mixture | 42.5 [37.5, 47.5] | A |
| | PRAZ | Mixture | 15.0 [10.0, 20.0] | F |
| NHDC | TAF | Pre-rinse | 39.5 [34.4, 44.5] | AB |
| | PRAZ | Pre-rinse | 29.2 [24.1, 34.2] | CD |
| ROSI | TAF | Pre-rinse | 40.7 [35.7, 45.7] | A |
| | PRAZ | Pre-rinse | 24.2 [19.1, 29.3] | CDE |

*[a]: Different letters show significant differences in mean intensity between samples.*

**Table S2**. Bitterness intensity of drugs and the mixtures with bitter blockers rated by the genetically informative panel (N=158) on a 100-point generalized visual analog scale.

| Blocker | Ligand | Bitterness intensity Mean [CI 95%] | Statistical grouping[a] |
|---|---|---|---|
| - | TAF | 68.1 [63.4, 72.8] | A |
| - | PRAZ | 46.2 [41.5, 50.9] | BC |
| - | PROP | 32.0 [27.3, 36.7] | D |
| NHDC | TAF | 43.1 [38.4, 47.7] | C |
| | PRAZ | 28.4 [23.7, 33.1] | D |
| ROSI | TAF | 54.3 [49.6, 59.0] | B |
| | PRAZ | 47.7 [43.1, 52.4] | BC |
| Simpler TZD | TAF | 68.6 [63.9, 73.3] | A |
| | PRAZ | 46.0 [41.4, 50.7] | BC |

*[a]: Different letters show significant differences in mean intensity between samples. All results with bitter blockers reflect the blocker given as a mixture with the bitter drug.*

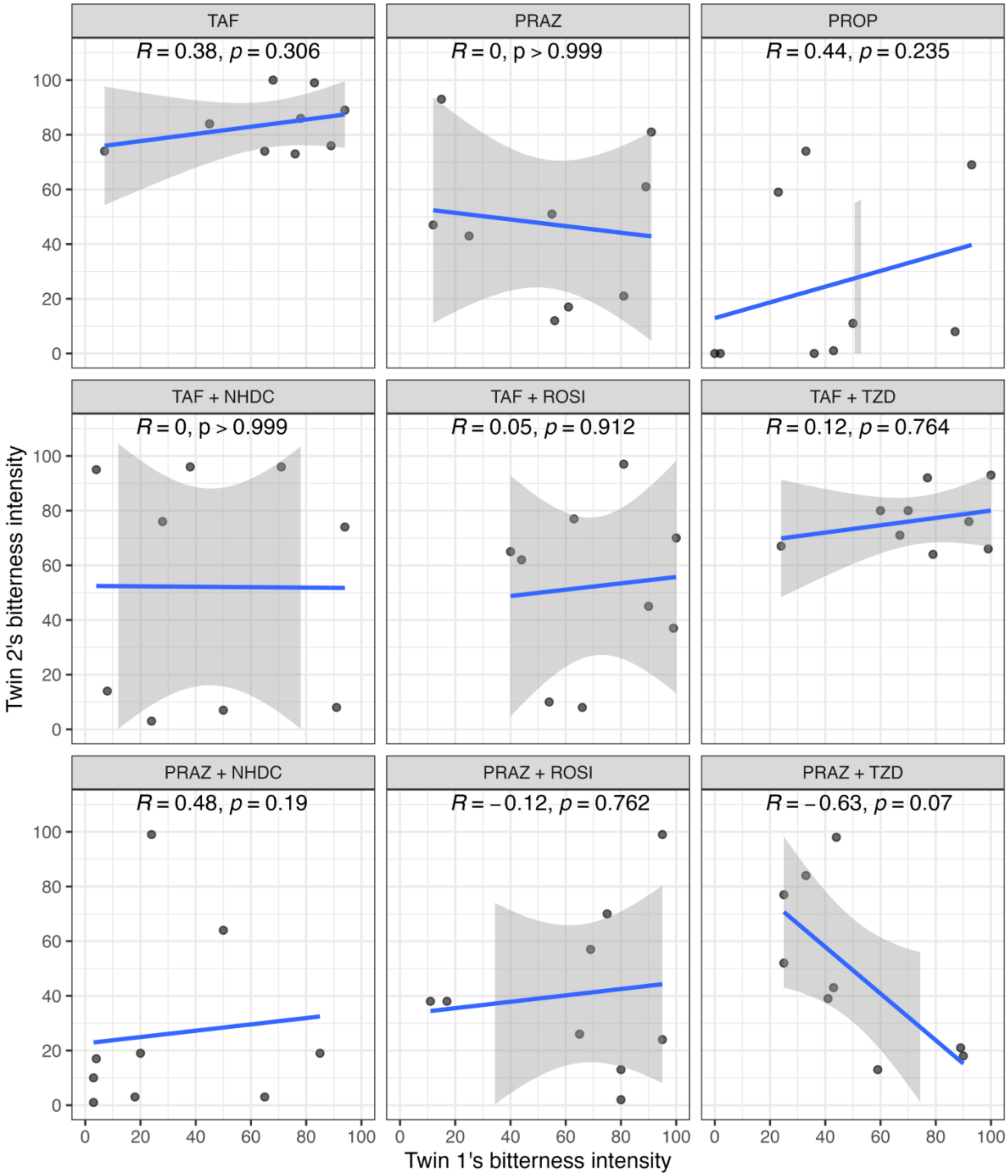


**Figure S1.** Spearman's correlation in bitterness intensity ratings between twin 1 and twin 2 for dizygotic twin pairs in the genetically informative panel (N=9). For abbreviations, see Table 1.

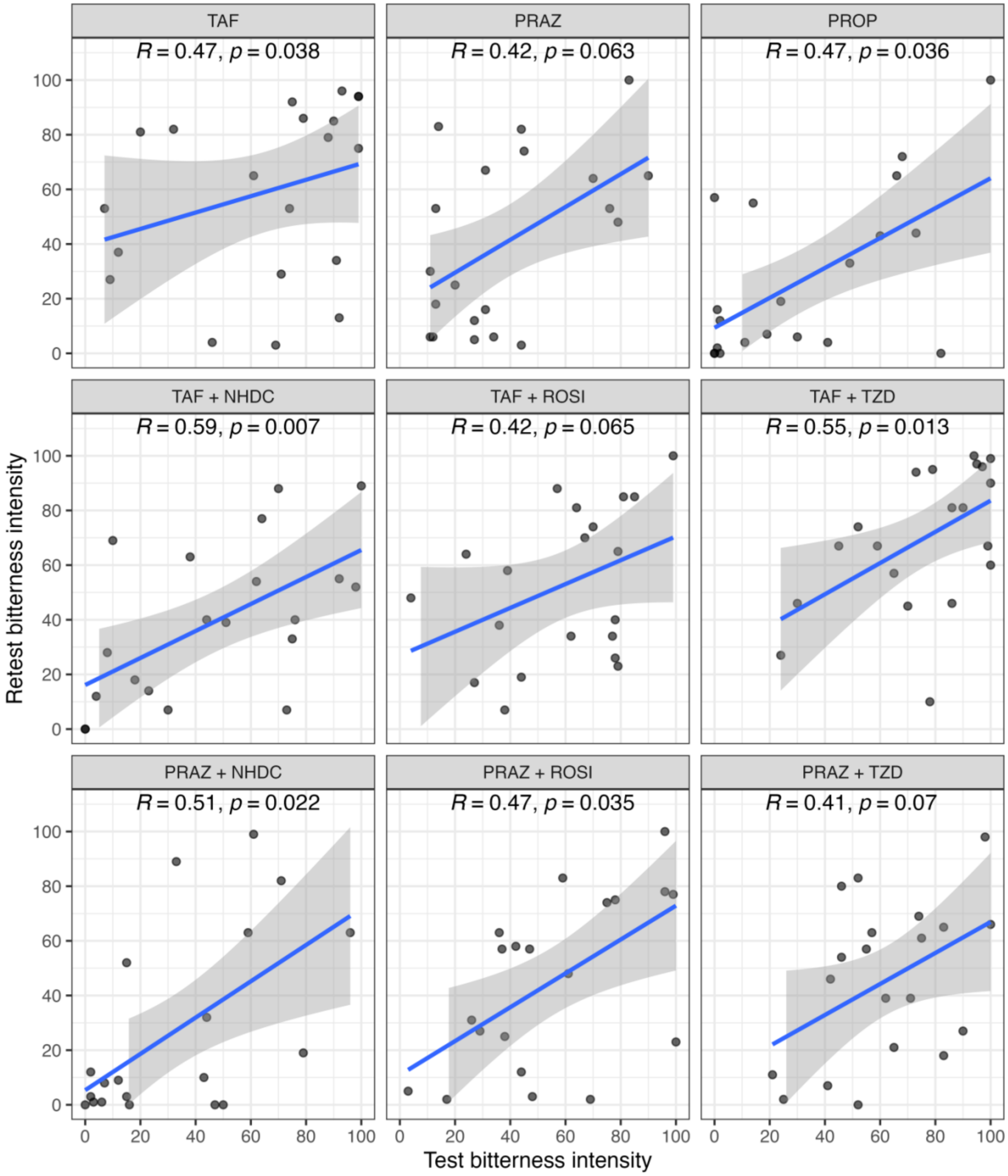


**Figure S2**. Spearman's correlation in bitterness intensity ratings between test and retest samples on different sessions in the genetically informative panel (n=20). For abbreviations, see Table 1.
